# When more is less: Dual phosphorylation protects signaling off-state against overexpression

**DOI:** 10.1101/295899

**Authors:** Franziska Witzel, Nils Blüthgen

## Abstract

Kinases in signaling pathways are commonly activated by multisite phosphorylation. For example, the mitogen-activated protein kinase Erk is activated by its kinase Mek by two consecutive phosphorylations within its activation loop. In this article, we use kinetic models to study how the activation of Erk is coupled to its abundance. Intuitively, Erk activity should rise with increasing amounts of Erk protein. However, a mathematical model shows that the signaling off-state is robust to increasing amounts of Erk, and Erk activity may even decline with increasing amounts of Erk. This counter-intuitive, bell-shaped response of Erk activity to increasing amounts of Erk arises from the competition of the unmodified and single phosphorylated form of Erk for access to its kinase Mek. This shows that phosphorylation cycles can contain an intrinsic robustness mechanism that protects signaling from aberrant activation e.g. by gene expression noise or kinase overexpression following gene duplication events in diseases like cancer.

## INTRODUCTION

The MAPK signaling pathway is one of the best studied signaling pathways due to its role in cell fate decisions like proliferation, migration and apoptosis and its critical role in development. Growth factors activate a receptor localised to the cell membrane, from where the signal is relayed by a cascade of kinases that activate each other by (reversible) phosphorylation on multiple sites. The terminal kinase, Erk, activates hundreds of cytoplasmic and nuclear targets (1). The activation of transcription factors induces a transcriptional response which ultimately manifests the cell fate decision.

An understanding of how such kinase cascades operate dynamically and quantitatively has been gained through a number of theoretical and experimental investigations. An early theoretical study showed that a single phosphorylation cycle can create a switch-like response (2). Later on, it was shown that the switch-like stimulus response profile of MAPK activity in *Xenopus* oocytes (3) can be explained by the *in vitro* distributive two-step activation mechanism of Erk (4). The mathematical description of phosphorylation cycles has its unique challenges as, opposed to metabolic networks, enzymes and substrates, all being kinases, mostly occur in similar concentrations. General concepts for modelling multisite phosphorylation (5–7) and for the analysis of multistability of these systems have been provided (8–10). Many studies focused on the stimulus-response relationship of a kinase that is activated by multisite-phosphorylation. The profile can be graded, biphasic, switch-like or bistable depending on a multitude of factors like the order (11) and/or processitivity (12, 13) of multisite phosphorylation, competition effects between modifying enzymes (5) or the sequestration of components within enzyme-substrate complexes (14–17). Some of the effects of competition and sequestration have been shown experimentally *in vivo.* For instance, the activity of Erk depends on the expression level of its substrates, as deactivating phosphatases and Erk substrates compete for access to Erk in *Drosophila* (18).

Next to the ability to process all-or-none decisions, signaling pathways should provide their response in a robust fashion: the signaling off-state needs to be robust to fluctuating levels of signaling pathway components and to transient weak signals (19, 20). Negative feedbacks are common in MAPK signaling and can provide robustness to Erk activity at various expression levels of Erk (21). However, some robustness might emerge from the phosphorylation cycle motif alone, as e.g. the amount of modified substrate approaches a limit for increasing levels of the substrate in a single modification cycle in its basal state (22).

Here we present a new mechanism that leads to robust stationary Erk activity at Erk overexpression, which emerges from the distributive kinetics of Erk phosphorylation. We find that for low pathway activity and increasing levels of total Erk, the stationary amount of active dual phosphorylated Erk shows a bell-shaped response: With increasing amounts of Erk, Erk activity increases until it reaches a maximum after which active Erk starts to decrease and eventually approaches zero. This bell-shaped response is due to the gradual saturation of Mek with its substrate and the subsequent competition of unmodified and single phosphorylated Erk for access to Mek. This response can be seen regardless of the order of Erk (de)activation and the kind of phosphatases involved in dephosphorylation of threonine and tyrosine on Erk. We derive an analytical approximation of the maximum in the bell-shaped response which allows to estimate the biological relevance of the phenomenon based on the catalytic rate constants.

Overexpression of signaling proteins is a common consequence of the massive genomic alterations in cancer and it is generally believed that this alteration will increase pathway activity or may cause spontaneous pathway activation. However, our results show that a distributive two-step activation of Erk has the potential to suppress excessive Erk activity and thus protects the signaling off-state against Erk overexpression, which may explain why Erk overexpression is rarely seen in tumors (23–25).

## MATERIALS AND METHODS

### Ordinary differential equation models

#### Basic model of Erk (de)activation

We model the 2-step activation and deactivation of Erk by assuming that the kinase and phosphatase forms a complex with its substrate in a reversible fashion (association rate constants *k*_onx_, dissociation rate constants *k*_offx_). (De)phosphorylation and release of the phosphatase/kinase from their respective modified substrates is assumed to proceed as one irreversible step with rate constant *k*_catx_. Within the index of kinetic rate constants x∈ {1,2} indicates the phosphorylation reaction in phosphorylation cycle 1 or 2, x∈ {p1,p2} the dephosphorylation the reaction in cycle 1 or 2. We denote the total concentration of active kinase ppMek as K_T_, the total concentration of phosphatase as P_T_ and the total concentration of Erk as Erk_T_. Complexes of kinase/phosphatase with their substrates are named C_x_/D_x_ where x∈ {1,2} indicates the 1st and 2nd phosphorylation cycle, see also the pathway scheme in Fig. 1A. The following ODE system describes the kinetics of its components:

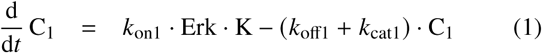

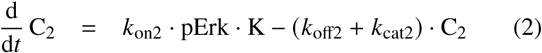

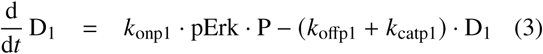

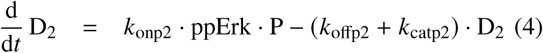

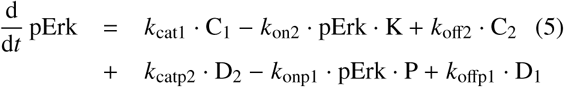

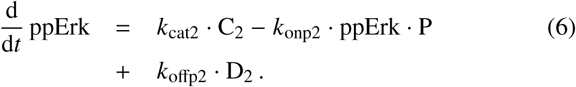

**Figure 1:**
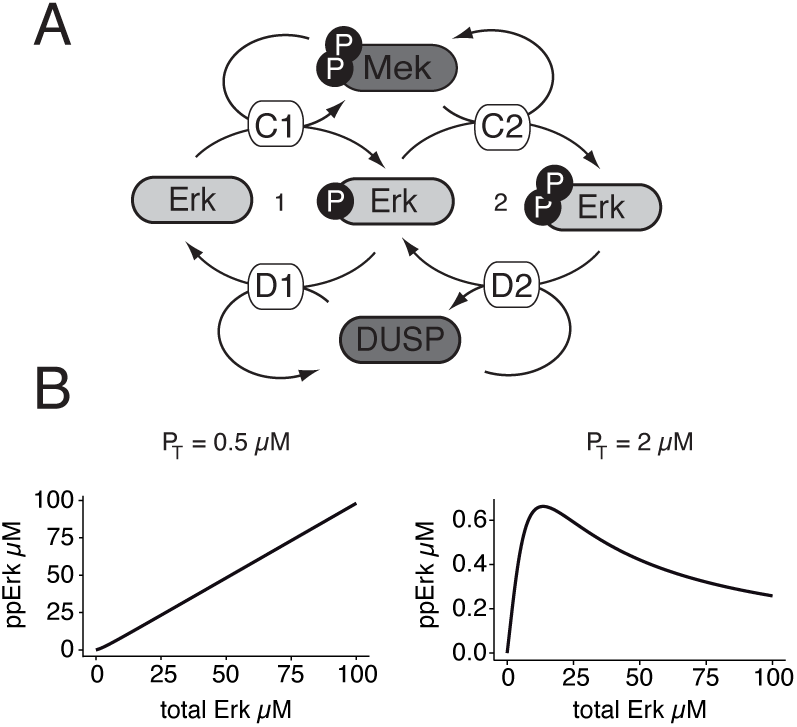
Bell-shaped response of active Erk as function of total Erk. A, Distributive (basic) model of Erk (de)phosphorylation. Enzyme-substrate complexes C/D_1/2_ are formed in a reversible fashion. DUSP = dual-specificity phosphatase. B, Simulation of stationary ppErk versus level of total Erk using the basic model for high (left) and low (right) pathway activity. Total amount of active Mek equals K_T_ = 1.2*μ*M. The amount of phosphatase has been chosen arbitrarily and is indicated with P_T_ at the top of the respective panel. All other parameters set as shown in table 1.

The concentrations of Erk, kinase K and phosphatase P can be calculated from the conservation relations:

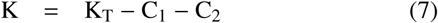

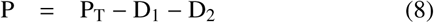

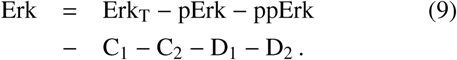

The kinetic parameters used for numerical simulation are shown in table 1. Several parameters of the model have been estimated *in vivo* in HeLa cells (26). All rate constants that describe the formation of an enzyme-substrate complex have been assumed to be identical, the same was assumed for the dissociation rates of these complexes.

**Table 1:** Table of parameters used in the basic model and in the model with two different phosphatases. Dephosphorylation rates *d*_1/2_ are used in the simplified model where we assume mass-action kinetics for Erk deactivation. *Measured apparent rates *r* = *k*_cat_/*K*_M_ were used to derive the catalytic rates according to the equation 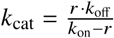
.

| param. [$s^{-1}\mu M^{-1}$] | value | comment |
| --- | --- | --- |
| $k_{on1}$ | 0.18 | measured in (26) |
| $k_{on2}, k_{onp1}, k_{onp2}$ | 0.18 | as $k_{on1}$ |
| $k_{cat1}/K_{M1}$ | $3.9 \cdot 10^{-2}$ | measured in (26) |
| $k_{cat2}/K_{M2}$ | $2.1 \cdot 10^{-2}$ | measured in (26) |
| param. [$s^{-1}$] | value | comment |
| $d_1$ | $6.7 \cdot 10^{-3}$ | $pYErk \rightarrow Erk$ (26) |
| $d_2$ | $4.0 \cdot 10^{-3}$ | $pTpYErk \rightarrow pYErk$ (26) |
| $k_{off1}$ | 0.27 | measured in (26) |
| $k_{off2}, k_{offp1}, k_{offp2}$ | 0.27 | as $k_{off1}$ |
| $k_{cat1}$ | $7.47 \cdot 10^{-2}$ | *calculated from (26) |
| $k_{cat2}$ | $3.57 \cdot 10^{-2}$ | *calculated from (26) |
| $k_{catp1}$ | $5.85 \cdot 10^{-2}$ | *calculated from (26) |
| $k_{catp2}$ | $3.15 \cdot 10^{-2}$ | *calculated from (26) |
| param. [$\mu M$] | value | comment |
| Mek total | 1.2 | measured in (26) |
| Erk total | 0.74 | measured in (26) |

#### Model with Erk deactivation by two different phosphatases

We describe the model in terms of modifications to the basic model. The conservation relations for the total kinase K_T_ and the total amount of Erk, Erk_T_, remain unchanged, however, we have to replace equation (8) by two equations for the conservation relations for one phosphatase, P_1_ and the 2nd phosphatase, P_2_

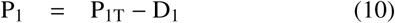

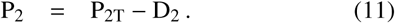

Model equations (1) and (2) remain unchanged. In equations (3) and (5) variable P is replaced by P_1_, in equations (4) and (6) variable P is replaced by P_2_. Kinetic parameters remain unchanged and can be found in table 1.

#### Ordered Model of Erk (de)activation

In this model we consider the two different forms of single phosphorylated Erk, pYErk (phosphorylated on tyrosine) and pTErk (phosphorylated on threonine). We model that Erk is phosphorylated and dephosphorylated on tyrosine first. Just like in the basic model we assume that binding of enzyme and substrate is reversible with rates *k*_onx_/*k*_offx_. Here x∈ {c1, cy2, ct2, dy1, dt1, d2} identifies the enzyme-substrate complex involved, where C1/CY2/CT2 is the complex of activating kinase with Erk/pYErk/pTErk and DY1/DT1/D2 the complex of phosphatase and pYErk/pTErk/pYpTErk. See also the pathway scheme in Fig. 6A. For qualitative analysis of this model we set the values of all kinetic parameters and of the kinase/phosphatase concentration to 1, unless stated otherwise. The following ODEs describe all components in this model:

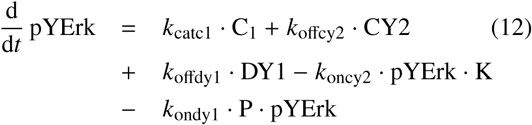

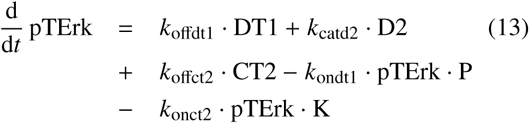

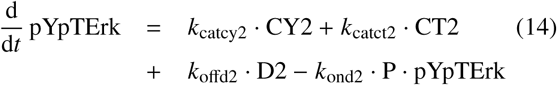

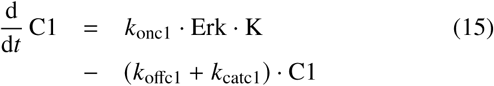

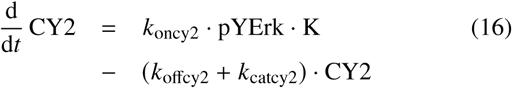

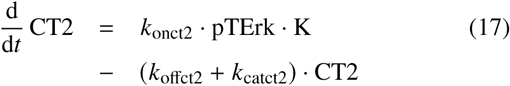

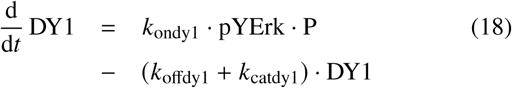

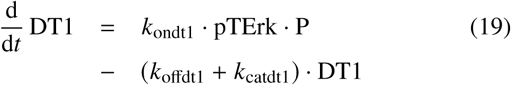

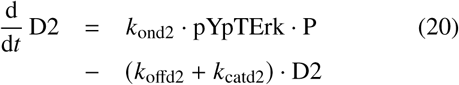

where the concentrations of Erk, kinase K and phosphatase P are given by the conservation relations

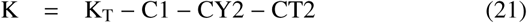

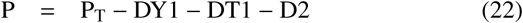

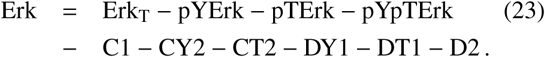

### Numerical simulations and calculations

All numerical simulations were carried out using MATLAB R2013b. To determine the steady state phosphorylation levels, the ODE system was solved by numerical integration (using the solver ode23s) until a time point where the solution approaches an equilibrium. Using the numerical root finding routine fsolve, the steady state was confirmed. Uniqueness of the steady-state was checked by starting from two opposing initial conditions, where either no Erk was phosphorylated initially, or all Erk dual phosphorylated. All analytical calculations have been verified using Wolfram Mathematica 8.

## RESULTS AND DISCUSSION

### Mechanistic model predicts reduced Erk activity at high Erk expression levels

To investigate the effect of changing concentrations of the target in a covalent modification cycle, we chose to model the activation of Erk. Erk needs to be phosphorylated on threonine and tyrosine within the TEY motif to be fully active (27). The only enzyme that catalyzes these two phosphorylation steps is Mek1/2. *In vitro* it has been shown that Mek cannot catalyze these two phosphorylations in one reaction (as processive enzymes do), but Mek preferentially phosphorylates Erk on tyrosine first (4, 28), and then the enzyme substrate complex dissociates and reforms for the second phosphorylation step (distributive mechanism) (29).

Erk is dephosphorylated and thereby inactivated by different types of phosphatases. Ubiquitous phosphotyrosine phosphatases like PTP remove the phosphorylation on tyrosine. DUSPs remove phosphates on both threonine and tyrosine (30). Another special characteristic of DUSPs is their specific localisation either to the nucleus or cytoplasm and their regulation by MAPKs themselves. Dephosphorylation by DUSPs is believed to follow a distributive scheme as well (31).

The direct proof for distributive kinetics has been provided by *in vitro* studies (28, 29). But a distributive mechanism has the potential to be converted to a quasi-processive one *in vivo.* Either molecular crowding (26, 32) or the anchoring to molecular scaffolds could increase the stability of the Mek-pErk complex and/or enable rapid rebinding of the latter. However, it has been shown that in mouse embryonic fibroblasts only the scaffold KSR and Mek1/2 form rather stable complexes in the cytoplasm, whereas the interaction of the scaffold with Raf and Erk is highly dynamic (33). Up to now the experimental evidence for distributive Erk phosphorylation *in vivo* outweigh the evidence for a quasi-processive mechanism (34–37).

We therefore developed a kinetic model which accounts for the (reversible) binding of Mek to Erk, its phosphorylation, and the (reversible) binding of DUSPs to Erk with subsequent dephosphorylation (see scheme in Fig. 1A). Phosphorylation and dephosphorylation were assumed to follow a distributive scheme. The ordinary differential equations (ODEs) and kinetic parameters that describe the kinetics associated with the presented reaction scheme can be found in Materials and Methods.

We then performed numerical simulations of the model, where we varied the total concentration of Erk. We noticed that the change of ppErk (dual phosphorylated Erk) upon increase of total Erk is qualitatively different for different activity ratios of the modifying kinase and phosphatase. For low concentration of the phosphatase, such as at P_T_=0.5 *μ*M (see Fig. 1B), when the maximal turnover rate of the kinase *v*_max_,_K_ = *k*_cat_,_K_ · K_T_ exceeds the maximal turnover rate of the phosphatase, ppErk rises linearly with total Erk. However, when the phosphatase dominates with P_T_=2 *μ*M, ppErk shows a nonlinear, bell-shaped dependence on total Erk (Fig. 1B). While ppErk increases first, it reaches a maximum and subsequently decreases for higher levels of Erk.

Puzzled by this non-intuitive behavior, we inspected how the different forms of Erk and its complexes with kinases or phosphatases change when the total amount of Erk is increasing. The single phosphorylated Erk increases monotonically with the Erk expression level, however, it approaches a limit (Fig. 2B). The ppMek-Erk enzyme substrate complex C_1_ shows a similar behavior as it approaches the concentration of total ppMek, here called K_T_ (Fig. 2B). This shows that ppMek becomes saturated with unphosphorylated Erk at increasing levels of the latter. This is reminiscent of a mechanism described previously as kinetic tumor supression for a single modification cycle. (22), and this mechanism will be key to understand the bell-shaped response of dual phosphorylated Erk, as shown below.

**Figure 2:**
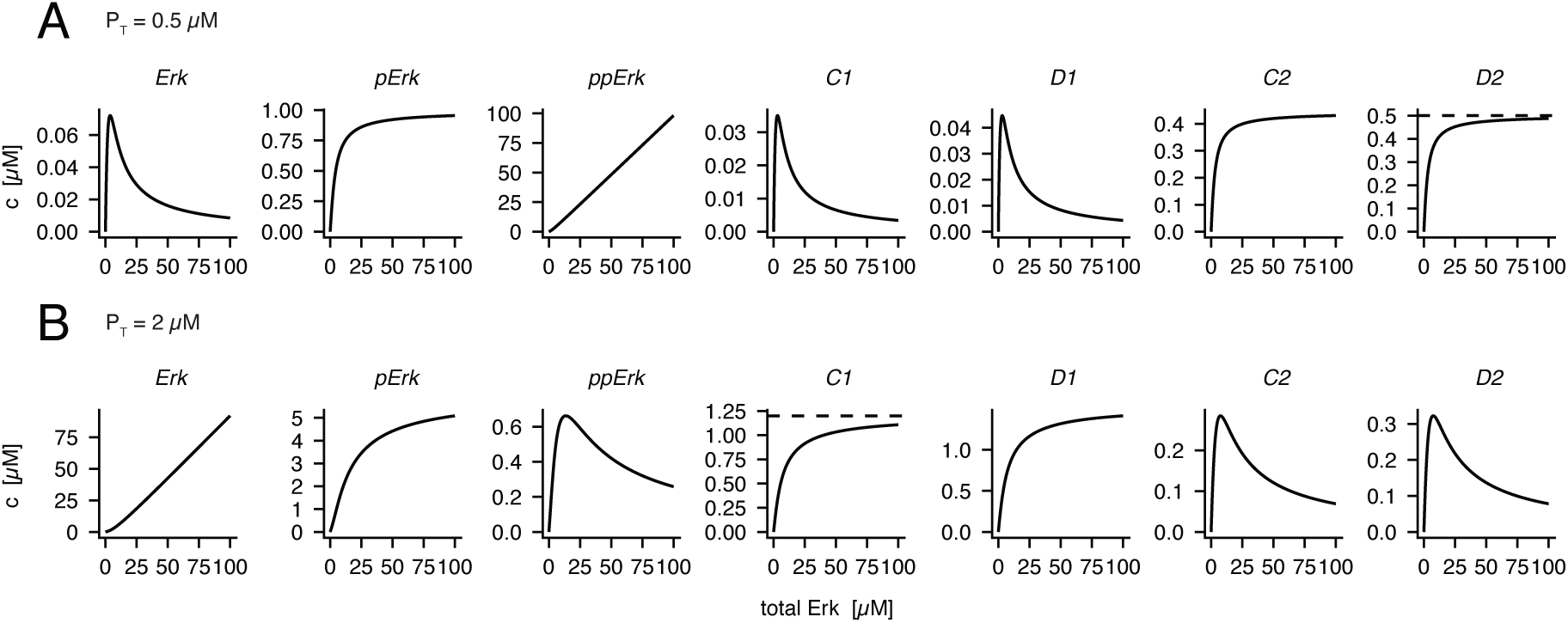
Steady state of the dual phosphorylation cycle when varying total amount of Erk. Simulation of the basic model with a total phosphatase concentration set to P_T_=0.5 *μ*M in A and P_T_=2 *μ*M in B. All other kinetic parameters set as listed in table 1. Dashed lines indicate the total concentration of the phosphatase in A and of the kinase in B.

### Limited activation in a single phosphorylation cycle

For now, let us assume that Erk is activated by a single phosphorylation that is provided by a kinase and removed by a phosphatase. Then, at low pathway activity, the amount of activated Erk has an upper limit (22). As the steady state of a single phosphorylation cycle has an analytical solution (2), this upper limit can be derived by calculating the mathematical limit of pErk as total Erk approaches infinity (22). However, there is an easier approach. As we consider a scenario involving large amounts of total Erk, we can assume Michaelis-Menten kinetics for the modifying enzymes, so the velocity of kinase/phosphatase is determined by its affinity to the substrate, *K*_M_,_K/P_, and its maximum turnover rate *v*_max_,_K/P_ (see Fig. 3). At low pathway activity *v*_max_,_P_ is larger than *v*_max_,_K_. As we consider a phosphorylation cycle, the velocities of kinase and phosphatase have to be identical in steady state (indicated by the black horizontal lines in Fig. 3). In consequence, the amount of pErk will be significantly smaller than the amount of unmodified Erk, as shown in Fig. 3A. If the level of total Erk is increased further, both enzymes are pushed to higher velocities, but the smaller *v*_max_,_K_ sets an upper limit to this steady state velocity (see Fig. 3B). In consequence, unmodified Erk accumulates while pErk approaches an upper limit. This limit can be derived from the steady state condition when the kinase operates at saturation:

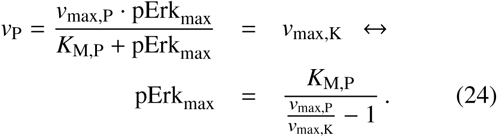

**Figure 3:**
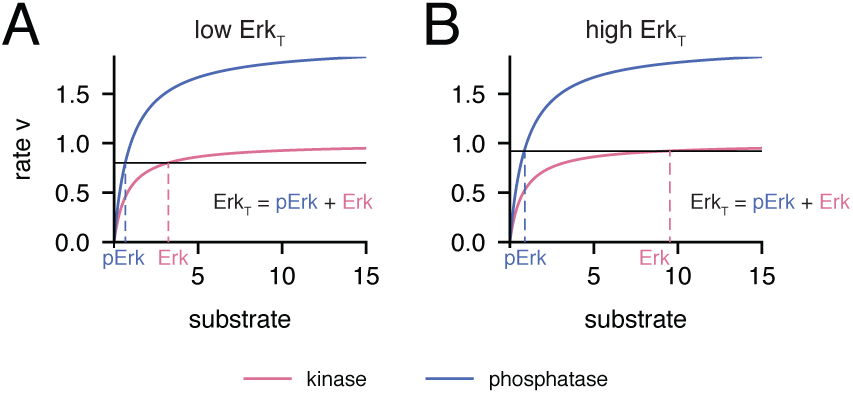
Overexpression insensitivity in a single phosphorylation cycle. The velocity of the kinase (pink) and of the phosphatase (blue) are shown as a function of substrate level according to Michaelis-Menten, where *v*_max_,_K_ < *v*_max_,_P_. In steady state, the velocity of the kinase equals the velocity of the phosphatase, which is indicated by the black horizontal line. The amount of substrates (Erk and pErk) follows as indicated by the dashed lines. A, for low amounts of total Erk, B, for high amounts of total Erk.

We see from equation (24) that the activity ratio of kinase and phosphatase directly influences the stationary level of phosphorylated Erk. When the maximal turnover rate of the phosphatase is twice the maximal turnover rate of the kinase, the maximal amount of phosphorylated Erk complys to the Michaelis-Menten constant of the phosphatase. The role of the phosphatases’ *K*_M_ is intuitive, as a weaker affinity of the phosphatase helps to pile up more of the activated species pErk.

It is now clear that when Erk is overexpressed the formation of active Erk is limited, because the kinase saturates and the phosphatase does not. The only parametric prerequisite for this effect is a lower *v*_max_ of the kinase compared to the phosphatase.

### The signal is attenuated further in a dual phosphorylation cycle

Also in the dual phosphorylation cycle the stationary level of single phosphorylated Erk rises with the total amount of Erk and finally approaches a limit, given that *v*_max_,_K_ < *v*_max_,_P_ (Fig. 2). This is due to progressing saturation of ppMek - however, now ppMek can either be bound in a complex with Erk (C_1_) or pErk (C_2_). As C_2_ approaches 0 and C_1_ approaches K_T_, (see Fig. 2B) active Mek apparently becomes sequestered within the first phosphorylation cycle. That means, two mechanisms shape the basal steady state amount of ppErk at Erk overexpression: saturation of ppMek and sequestration of ppMek in the first phosphorylation step. Consequently, the phosphatase is also drawn into the first phosphorylation cycle - complexes D_2_ and C_2_ decrease for rising levels of total Erk (Fig. 2B).

When the condition is reversed, so when *v*_max_,_K_ > *v*_max_,_P_, all intermediate species of the dual phosphorylation cycle behave in a mirror-inverted fashion, e.g. unphosphorylated Erk exchanges its concentration profile with the profile of dual phosphorylated Erk. The phosphatase saturates in the 2nd phosphorylation cycle and draws most of the kinase into the 2nd cycle (Fig. 2A).

As either the kinase (phosphatase) is sequestered in the first (second) phosphorylation cycle, the limit of single phosphorylated Erk in a dual phosphorylation cycle can be calculated like in a single phosphorylation cycle:

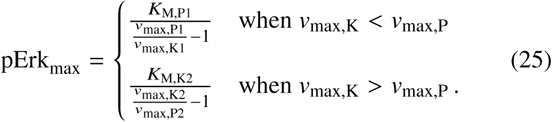

*K*_M,P1_ refers to the affinity of the phosphatase in cycle 1, which is its affinity to pErk. Likewise *K*_m_,_K2_ refers to the affinity of the kinase in cycle 2 – the affinity of the kinase to pErk. Maximum turnover rates *v*_max_ are labelled accordingly. Figure 4A shows the amount of single phosphorylated Erk in a dual phosphorylation cycle for increasing amounts of Erk and the calculated limits using the equation (25).

**Figure 4:**
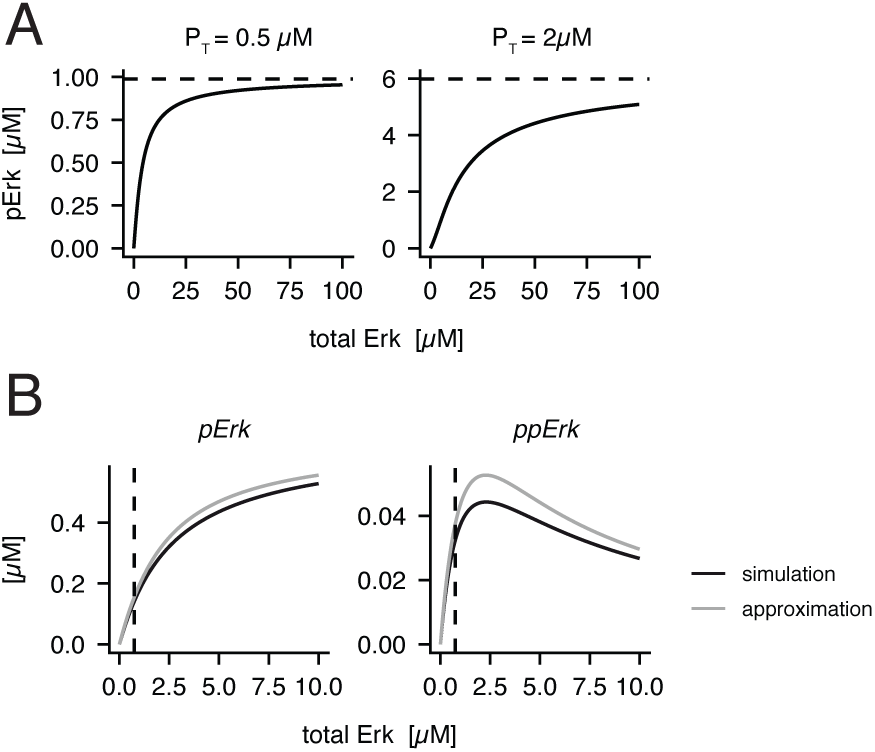
Quantification of Erk activation limit. A, Numerical simulation of the amount of pErk in a dual phosphorylation cycle according to the basic model (black line) for high (left) and low (right) pathway activity. The analytical limit of pErk (see eq. (25)) is indicated by the dashed line. All parameters chosen as listed in table 1. B, The steady state level of pErk and ppErk at varying levels of Erk_T_ was simulated with the simplified model that features distributive dual phosphorylation of Erk by Mek and mass action rates of dephosphorylation. The conservation relation for K_T_ and Erk_T_ is either exact (simulation, black line) or approximated according to eq. (28) and (29) (analytical approximation, gray line). The dashed line indicates the concentration of Erk in HeLa cells (26).

### A simplified model explains limited activation in a dual phosphorylation cycle

To improve our understanding of how the various rate constants shape the maximum of Erk activation in a dual phosphorylation cycle we sought to simplify our basic ODE model (1)-(6) in a way that will allow us to calculate a closed form of the steady state. Limited activation of Erk is seen when ppMek is shared between two cycles and eventually saturates and sequesters in one of the cycles. The phosphatases keep working far from saturation, so that we can model their catalysis with mass-action kinetics instead. Thus the model equations (1) and (2) remain unchanged but the equations (3) and (4) that describe the temporal development of the phosphatase in complex with its two different substrates can be dropped. Assuming that dephosphorylation of single phosphorylated Erk proceeds with rate *d*_1_ and dephosphorylation of dual phosphorylated Erk with rate *d*_2_, equations (5) and (6) are rewritten to

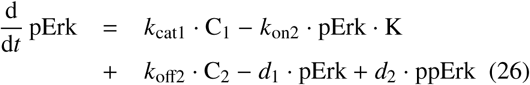

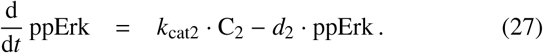

Even with this modification, the explicit description of all components in steady state is impossible, which is generally true when the various enzyme-substrate complexes are appreciable compared to the concentration of free substrate and product (38). However, we can approximate

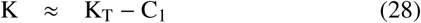

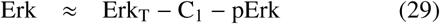

because the concentration of the complex formed by ppMek and monophosphorylated Erk, C_2_, is significantly smaller than C_1_ and ppErk has the smallest contribution to the total level of Erk.

In equation (28) and (29) Erk_T_ and K_T_ denote the respective total enzyme concentrations of Erk and ppMek. The steady state of this simplified system has a closed form and reads:

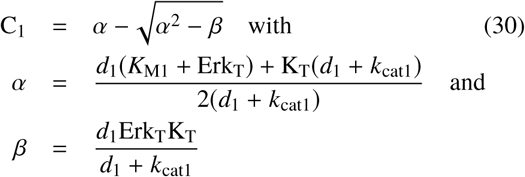

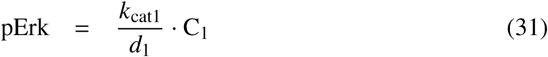

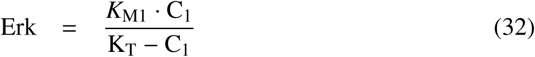

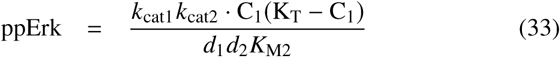

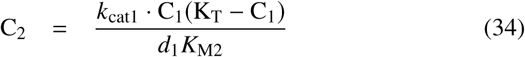

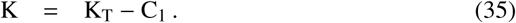

Here, *K*_M1/2_ refers to the Michaelis-Menten constant of the kinase in the first/second phosphorylation cycle. The approximation of the steady state captures the correlation of phosphorylated Erk and total Erk qualitatively as well as the order of magnitude in phosphorylation, as can be seen in a direct comparison of the numerical solution of the system with mass-action kinetics for dephosphorylation with the analytical approximation (Fig. 4B) where the conservation relations of Erk and ppMek have been truncated as shown in equation (28) and (29).

Using the analytical solution from equation (33) we can now derive the concentration of Erk at which ppErk is maximal. The derivative of ppErk by the level of total Erk

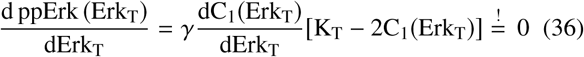

equals zero at the maximum with

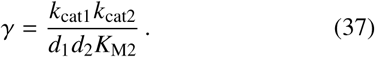

Condition (36) is only fulfilled when

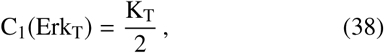

as C_1_ grows with the amount of Erk_T_ until saturation of the kinase with Erk, the first factor, 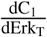, is never zero. The level of total Erk in the cell leading to maximal activation is the one where half of the total available kinase ppMek is sequestered in a complex with unphosphorylated Erk. Condition (38) allows for the exact calculation of the maximum coordinate to

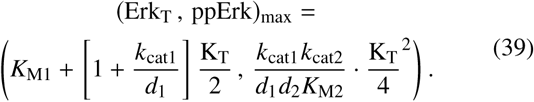

Note that this model will always create a bell-shaped response as it is built on the assumption that the phosphatases cannot saturate - changing any parameter in this model will on;y alter the position and/or height of the peak of activation. The maximal ppErk level is proportional to the square of the kinase concentration which reflects the two step nature of the activation process. A higher affinity of the kinase to un-phosphorylated Erk (smaller *K*_M1_) enforces sequestration and thus shifts the position of the peak to smaller levels of Erk. A higher affinity in catalysis of the 2nd phosphorylation (smaller *K*_M2_) increases the activation level. Only the catalytic rates of the 1st modification cycle (*d*_1_ and *K*_cat1_) influence the peak position, which suggests that the activity ratio of kinase and phosphatase in the cycle converting between Erk and pErk creates the prerequisite for limited activation.

In this model C_1_ approaches the level of K_T_ for increasing concentrations of Erk. It follows from equation (31) that the limit of single phosphorylated Erk amounts to

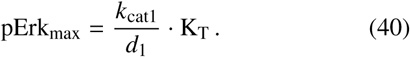

### Quantification of the activation limit

Using the equations (39) and (40) with the kinetic parameters measured in HeLa cells we can now estimate whether the kinetic suppression of excessive amounts of active Erk might play a role *in vivo.* Assuming that only 5% of the cellular Mek is activated, maximal levels of active Erk can be found at 2.3 *μ*M which is about 3 fold more than the average Erk expression level measured in HeLa cells (Fig. 4B). Also, only 2% of Erk is activated at the peak, which means that 5% Mek activity is attenuated to only 2% of Erk activation at the peak. For the physiological concentration of Erk, at 0.74 *μ*M, indicated with the dashed vertical line in Fig. 4B, the relative Erk activation is at 4.5%. Single phosphorylated Erk approaches a limit, which accords to 0.67 *μ*M.

With the help of the analytic equations derived here, the maximal activation level of a target can be estimated for any single or dual phosphorylation cycle, given that the catalytic rates are known. In case of Erk activation in HeLa cells, the mechanism which limits Erk activation is effective already at 3x overexpression, which can be considered mild in comparison to the observation that Erk concentrations vary about 3 fold between clonal cells (39).

### Different phosphatases can be involved in Erk deactivation

So far we have assumed that one enzyme is responsible for (de)phosphorylation of threonine and tyrosine on Erk. But dual-specificity phosphatases are a class of phosphatases whose expression is highly regulated in concentration and location (40). Under some circumstances they might not even be the main phosphatases responsible for deactivation of Erk. In the scenario where dephosphorylation of threonine and tyrosine is carried out by two different phosphatases, the activity ratio of kinase and phosphatase may differ in the two cycles. To test the prerequisite for the bell-shaped response under these circumstances we have adapted the basic model to include two different phosphatases as shown in the scheme in Fig. 5A.

**Figure 5:**
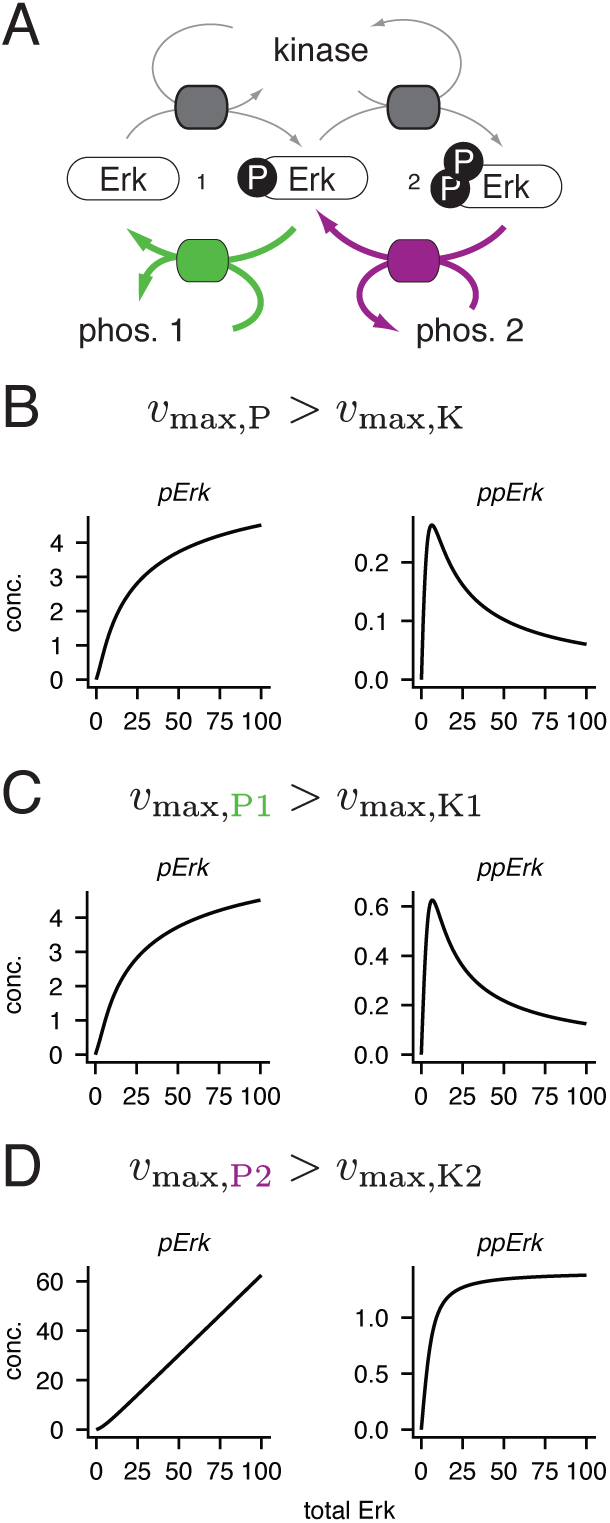
Limited activation in dual phosphorylation cycles where different phosphatases catalyse the first and second dephosphorylation. A, The basic model was modified to a scheme in which two different phosphatases deactivate Erk (see Material and Methods section). We show the steady state amounts of pErk and ppErk for different levels of total Erk when *v*_max_ of the phosphatase exceeds the level of *v*_max_ of the kinase in both cycles (B) and when the phosphatase has a higher maximum turnover rate than the kinase in only one out of the two cycles as indicated at the top of the panels C and D.

When the two phosphatases outcompete the kinase in both cycles, ppErk shows the same non-linear profile as was seen before (Fig. 5B). The bell-shaped response of ppErk is also found when phosphatase 1 has a larger turnover rate than the kinase, but not phosphatase 2 (Fig. 5C). However, if only the phosphatase 2 has a higher maximum turnover rate than the kinase, the formation of single phosphorylated Erk is proportional to the amount of available Erk and ppErk approaches a limit like in a single modification cycle (Fig. 5D).

It can be concluded that as long as the phosphatase dominates the activity of the kinase in at least one cycle, activation of Erk is limited even at higher expression levels. However, the model with two phosphatases clearly shows that a dominant activity of the phosphastase within the first phosphorylation cycle is sufficient for the bell-shaped profile of dual phosphorylated Erk.

### Prediction for the ordered model of Erk modification

Experimental evidence supports the hypothesis that the activating and deactivating modification of Erk proceeds in an ordered fashion: Erk is phosphorylated and dephosphorylated on tyrosine first. We have built a model to account for this by explicitly considering the 3 different states of phosphorylated Erk, pYErk, pTErk and pYpTErk. We assume that the conversion from Erk to pTErk as well as the conversion from pYpTErk to pYErk do not occur (see model equations in Material and Methods and a model scheme in Fig. 6A). From the results above we concluded that the bell-shaped response of active pYpTErk occurs only if the maximum turnover rate of the phosphatase exceeds the maximum turnover rate of the kinase within the first phosphorylation cycle. To test whether this condition still holds, we simulate the stationary amount of active Erk while varying the total amount of Erk with a parameter set in which the concentration of the kinase and the phosphatase equal 1 *μ*M and all other kinetic parameters are set to 1. Now the first phosphorylation cycle in the ordered scheme constitutes the cycle between Erk and pYErk. If we set the rate constant *k*_catdy1_ to 2 (while keeping all other parameters at 1), the condition for the bell-shaped pYpTErk is fulfilled. And indeed we find the previous saturation of pYErk to a limit value and a bell-shaped profile of pYpTErk (see Fig. 6B). In contrast, as pTErk is only created from pYpTErk in this ordered scheme, this species also shows a bell-shaped response curve. Here, the kinase is saturated in complex C1 and the phosphatase operates far from saturation, as described previously.

**Figure 6:**
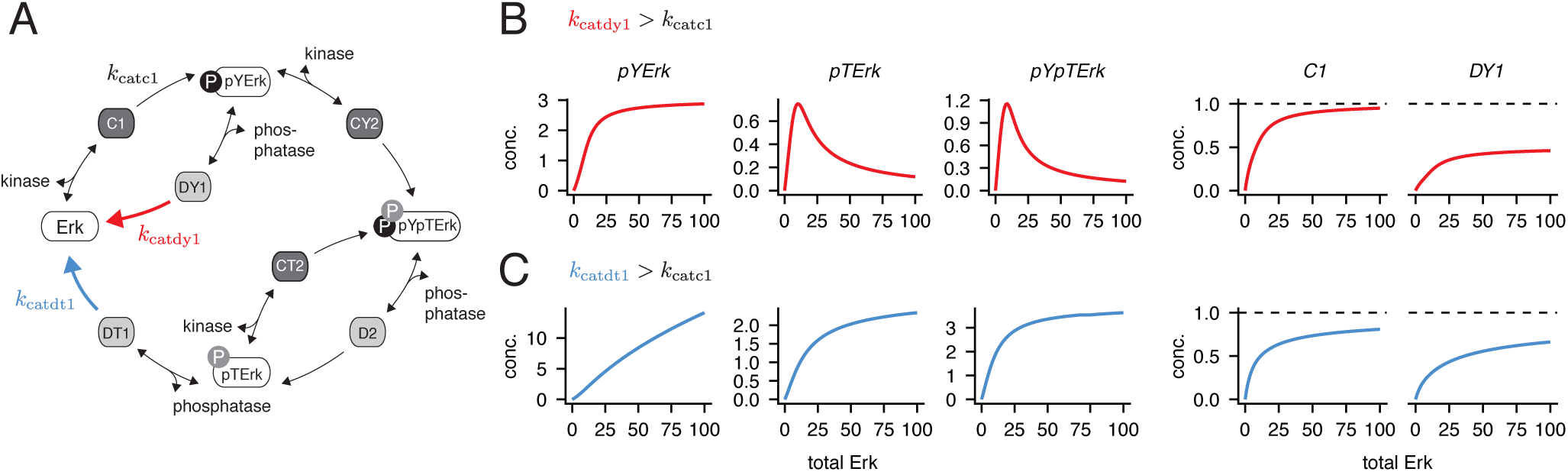
Limits to active Erk in the ordered model of Erk (de)activation. A, According to the ordered model Erk is phosphorylated and dephosphorylated on tyrosine first. B, Simulation of steady state of the different modification states of Erk and of the enzyme-substrate complexes C1 and DY1 when varying the amount of total Erk. All kinetic parameters and concentrations of modifying kinase and phosphatase have been set to 1, except for *k*_catdy1_ = 2. As a consequence *v*_max_ of pYErk dephosphorylation exceeds *v*_max_ of Erk phosphorylation to pYErk. C, Like in B, but now *k*_catdt1_ is the only parameter set to 2, which makes *v*_max_ of pTErk dephosphorylation larger than *v*_max_ of Erk phosphorylation to pYErk.

Alternatively, one can ask what happens when we assume that the dephosphorylation rate from pTErk exceeds the rate of phosphorylation from Erk to pYErk, by setting all parameters to 1 but the rate *k*_catdt1_ = 2 (Fig. 6C). Here, significant amounts of pYErk can be formed which serve as substrate to the second step of phosphorylation. In consequence, we see a plain limit to the amount of active pYpTErk, as would be the behaviour in a single modification cycle for high levels of substrate. Again, pTErk has the same concentration profile as pYpTErk, because it is only being created from it. Interestingly, both the kinase and the phosphatase are drawn into the first phosphorylation cycle here, i.e. the kinase is sequestered in complex C1 and the phosphatase in complex DY1.

We can conclude that we still find a bell-shaped pYpTErk response profile when the dephosphorylation rate of the tyrosine residue of Erk’s activation loop exceeds the phosphorylation rate of this residue. However, also when dephosphorylation of the threonine residue dominates the activating phosphorylation, we find robustness of the signaling off-state to increasing amounts of total Erk, as pYpTErk does not rise in a linear fashion, but approaches a limit.

## CONCLUSION

When the activity of a signaling protein is modified by the addition of one phospho-group, the signaling off-state is robust to increasing amounts of the protein itself, as the modifying kinase saturates eventually. If the activity of a protein is regulated by two consecutive phosphorylation events, the formation of dual phosphorylated active protein at increasing levels of protein is suppressed even further as the modifying kinase gets saturated with its substate and additionally gets sequestered within the first phosphorylation step, which makes it less available for catalysis of the second phosphorylation step. The prerequisite for this phenomenon is the distributive nature of two-step activation. As of now there is no clear consensus as to whether Erk is activated in a distributive fashion *in vivo.* However if so, the kinetic suppression of excessive amounts of active Erk described here in combination with the multitude of negative feedbacks present in MAPK signaling might explain why increasing expression of Erk alone would not confer a growth advantage to cells and why overexpression of Erk is rarely found in cancer in contrast to e.g. the frequent overexpression of receptors of the HER family.

## AUTHOR CONTRIBUTIONS

FW carried out all simulations and analytical calculations. NB and FW designed the research and wrote the article.

## ACKNOWLEDGMENTS

We thank the Federal Ministry of Education and Research for funding (grants MMML-Demonstrator, 031A428F and Map-Tor-Net, 031A426A).

